# A proposal for the quantum mechanical study of genomic mutations driven by environmental stressors

**DOI:** 10.1101/2024.05.24.595767

**Authors:** Héctor Mejía-Díaz, Diego Santiago-Alarcon, Salvador E. Venegas-Andraca

## Abstract

We propose a novel quantum statistical method to characterize DNA point mutations under the influence of UVC radiation, salinity, and temperature. We consider an open quantum system composed of an external environment — high-energy photons — coupled to the DNA molecule, and using energy considerations we estimate free parameters in a many-body Hamiltonian, to characterize critical behaviour in the system. The model presented here offers the advantage that one does not explicitly need to know each detail of the interaction between the base-pairs and the environment, by knowing whether the effect is to associate or dissociate the DNA molecule, one can incorporate it into the model.

We show that by fine-tuning the free parameters, the model gives results that are within a biologically realistic range in the energetic scale. Importantly, specific heat values show that by coupling DNA to an external bath, the system dynamics leads to larger fluctuations at slightly lower temperatures. Following this research strategy, one could use experimental data to extract a correlation between specific heat and DNA changes, which would provide insight to predicting evolutionary changes.

## 1 Introduction

During the last six decades, empirical evidence supporting non-genetic and non-random mutational processes in evolution has accumulated [1, 2]. Some genomic studies show certain patterns of induced genetic variation [3, 4], but the physical principles explaining why only certain regions of the genome are more susceptible to mutation is currently not understood. The recent development of Assembly Theory (AT) [5] has improved our understanding of selection and evolution, and helped us understand how life and, more generally, any complex object can come into existence from fundamental building blocks (e.g., molecular bonds). AT shows that from all the large universe of complex objects that can exist, only a highly restricted set is observed – what they called “assembly observed”; akin to historical and developmental constraints leading to a smaller evolutionary space of options. Also, AT shows how complex objects can develop through the interplay of selection and the assembly rules of the basic building blocks (i.e., physical constraints). Yet, once we have a complex object like the genome, AT still does not explain why evolution can occur rapidly within those constraints, why life can produce fast adaptive mutations (i.e., biased or non-random mutations). Thus, no current compelling theoretical framework from quantum physics (the appropriate scale of the genome) exists to help scientists understand the rationale for mutational biases and why adaptive evolution can occur rapidly even in complex organisms. Population genetics works with the observed standing variation in populations (the assembly space in AT), analyzes the change in frequency of variants (i.e., alleles, genotypes, number of objects with a specific assembly index in AT) across generations, and assumes a certain mutation rate (i.e., the rate of discovery and rate of production in AT). As its name implies, population genetics pertains to the population scale of organization and is thus a group level summary of what forces proceed below the population level in the genome, which is the biological scale where mutations or genomic variations are generated.

Quantum biology is a young discipline that seeks to explain key mechanisms of life using quantum physics [6], including evolution [7]. There is solid evidence demonstrating that the quantum mechanisms of superposition and entanglement are key for the occurrence of photosynthesis, and that enzymatic reactions are based on the proton tunneling mechanism, which is responsible for overcoming the energy barriers needed for a reaction to occur [8]. Yet, the work on quantum evolution has lagged behind because of a lack of an evolutionary foundation where the fundamental quantum physics can be applied (but see proposal in [7]). To appreciate why the quantum realm is relevant for evolutionary genomics, we must consider the following: (1) the DNA double strand is hold together by the sharing of hydrogen protons, (2) the regulatory and enzymatic processes occurring in the genome are based on quantum mechanisms, (3) the sequence of discrete nucleotides can be described as digital objects, where mutations represent quantum jumps between different states (i.e., nucleotides) in a DNA sequence that turns them into quantum digital objects where computations can be performed, and (4) the genome is a systems biology structure defined by interactions (e.g., network regulatory systems) that makes it a computational or cybernetic assembly [7]. Thus, to understand the genome we must study it with the computational methods at the appropriate scale of the quantum realm, which leads us to the development of quantum algorithms or quantum walks [QW] (e.g., [9, 10]) that represent the biological algorithms explaining the workings of the genome from fundamental physical principles.

The theory of random (spontaneous) mutations does not explain mutational biases due to the physical structure of the DNA [11]. In addition, we need to take into account quantum effects - such as proton tunneling - that will further increase biases in genome variation production due to its interaction with the environment. Adding an environment that interacts with the system under study has been shown to reconcile theory and experiment regarding hydrogen bond length in DNA [12]. Considering that genomic mutations are biased [2] and that the DNA molecule is a quantum system, such biases can be characterized by using quantum tools that consider the interactions between the physical environment (e.g., radiation) and the DNA molecule. In this sense, the mutations caused by such interactions require an exchange of fundamental particles [13]. The exchanged particles act as an external environment that determines the possible outcomes. Photons represent a suitable particle to understand the effect of the environment on the DNA molecule, particularly on the process of generating biased mutations. From physical principles, photons have no self-interactions and interact weakly with the air, which provides a system that is locally rich in information [6]. In addition, Ogryzko et al. considered that directed point mutations and deletions are a result of selecting possible outcomes created by an external environment, also asserting that mutational predictions must be of a statistical nature [14]. Thus, a quantum statistical analysis is a logical choice to include a broad set of elements into the study of mutations.

Here, we construct a quantum mechanical description of a synthetic gene interacting with a bosonic bath (UVC radiation). The gene is treated as a set of *N* base-pairs interacting with *M* bosonic modes (radiation frequencies) at different temperatures. We approximate the partition function of the system to extract thermodynamic properties of the system, namely its (average) energy, heat capacity (energy necessary to increase the temperature by 1 K), and specific heat (energy necessary to increase the temperature of 1 g of substance by 1 K). We follow this approach due to its statistical nature, considering that several contributions of energy can be categorized concisely into DNA self-energy, bosonic bath energy, and DNA-radiation energy. The operator representation of the energies is achieved in analogy with the well-known Ising and Dicke-Ising models — see for instance [15, 16, 17] —, frequently employed in condensed matter physics.

This article is organized as follows: firstly, we present the construction and the rationale for the model of gene-radiation interaction, where we give a small review of the relevant operators used in the model. We also identify them with the biological entities that they represent and show how to combine them into an effective Hamiltonian to describe a quantum open biological system. Secondly, we briefly describe how one can extract the thermodynamics of the model via the partition function, and what it means in our current biological context. Thirdly, we present the construction of a toy-model gene composed of a few base pair elements interacting with only a few bosonic modes, and provide evidence that the model is biologically realistic, in the sense that small interactions lead to small changes in the gene, while larger interactions result in important changes in the behavior of the DNA molecule. Thus, we provide a novel open quantum model to describe genomic mutations. This model can help to understanding why genome evolution can occur rapidly under the influence of some environmental factors, while also quantifying the DNA changes into thermodynamic quantities.

## 2 Gene-radiation interaction

We treat the problem of determining the direct and indirect effects of radiation at different temperatures on mutations, under the formalism of open quantum systems, — see, for instance [18] —, where the general form of the Hamiltonian operator^1^ is

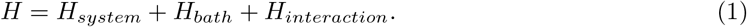

*H*_*system*_ is the Hamiltonian that determines the dynamics of a system of interest, whose physical properties we would like to know. Here, we set *H*_*system*_ = *H*_*DNA*_. The second term, *H*_*bath*_, refers to an external physical system, much larger than the one under study, that will affect its dynamics; here, *H*_*bath*_ refers to external UVC radiation. Finally, *H*_*interaction*_ couples the system and the bath.

### 2.1 Relevant Operators

The Hamiltonian under consideration consists of Pauli operators, to describe the DNA, and bosonic operators, to describe the bath. For completeness, we sketch a simple description of them, before giving the explicit form of the Hamiltonian.

#### 2.1.1 The Pauli operators

The Pauli operators, which we will denote by *σ*^*x*^, *σ*^*y*^ and *σ*^*z*^, satisfy the commutation relations

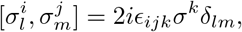

where *i, j, k* ∈ {*x, y, z*}, *δ*_*lm*_ is Kronecker’s delta,

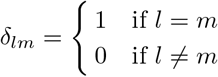

and *ϵ*_*ijk*_ is the Levi-Civita symbol,

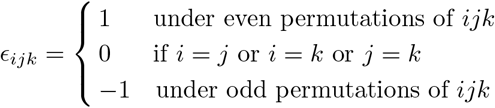

The subscripts *l* and *m* identify on which (Hilbert) space the Pauli operators are acting on; operators acting on different spaces always commute. If we denote the eigenstates of 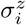 as |0⟩_*i*_ and |1⟩_*i*_, the action of *σ*^*x*^ and *σ*^*z*^ on these states is given by^2^

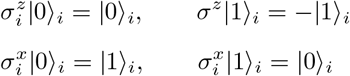

It is also relevant to notice that the Pauli operators are traceless,

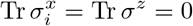

#### 2.1.2 The bosonic operators

From the formalism of second quantization (see [20], for example), we introduce the creation, annihilation, and number^3^ operators, denoted by *b*^*†*^, *b* and 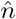, respectively. We use these operators to describe the bosonic bath and, combined with the Pauli operators, the couplings between the DNA and the bath.

The bosonic creation and annihilation operators are defined by the commutation relation

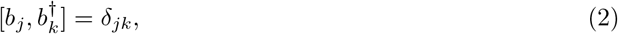

where the subscripts refer to the space in which each operator acts, as before. From the definition of the number operator, 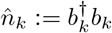, we get the following commutation relations

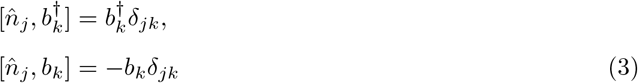

The action of the bosonic operators on the Fock states, |*n*⟩_*k*_, is

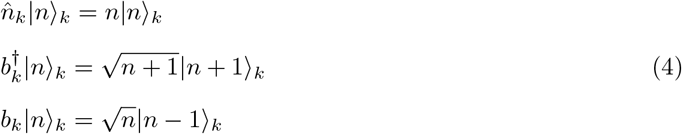

where a Fock state is a state with *n* particles (photons, in the present case).

### 2.2 DNA-Radiation Hamiltonian

We use the previously defined operators to propose a Hamiltonian operator that captures the dynamics of the interaction of DNA with external radiation. In eq. (1), we identify the following terms

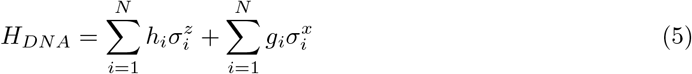

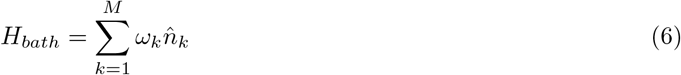

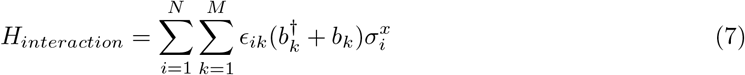

In *H*_*DNA*_, eq. (5), we to capture the dynamics of the DNA with two energetic contributions. The first term, dependent on 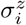, corresponds to the ground state energy of the unperturbed DNA molecule; each base pair is said to be in a given state with an energy *h*_*i*_, and upon summing on the *N* base pairs in the DNA chain, we get this total energy. The second term, dependent on 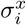, allows external elements — such as it being exposed to an external saline medium — to indirectly change the energy of the DNA molecule; indirect effects are tuned by the constants *g*_*i*_ (recall the effect of *σ*^*x*^ on the eigenstates of *σ*^*z*^).

The bath Hamiltonian, eq. (6), is composed of a combination of harmonic oscillators, where an additive constant has been omitted, and *ω*_*k*_ are the energies associated with each bosonic mode^4^. We consider *M* oscillators, each contributing to the total energy as an individual oscillator.

The interaction part of the model, eq. (7), based on the Ising-Dicke model [17, 16, 21, 22, 23, 15], models the interaction of the radiation and the DNA molecule. This term considers that, for a given bosonic mode, its coupling to each base pair in the molecule is the same, *ϵ*_*ik*_. By summing over all the possible interactions, this term drives energy fluctuations on the DNA.

### 2.3 Extracting information

To extract physical information from the system, we use the partition function from quantum statistical mechanics (for a review, see [24]); particularly, we use the grand partition function, where the system can exchange energy with the environment. The partition function is defined as

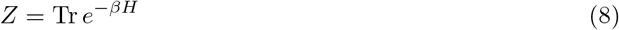

where 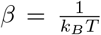, *k*_*B*_ is Boltzmann’s constant, and *T* is the (absolute) temperature. The partition function allows us to compute the energy, specific heat and heat capacity of the model, thus enabling us to predict most stable configurations under different external pressures at different temperatures.

After having computed the partition function, we can extract averages for the energy

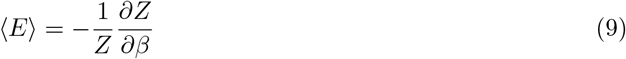

heat capacity,

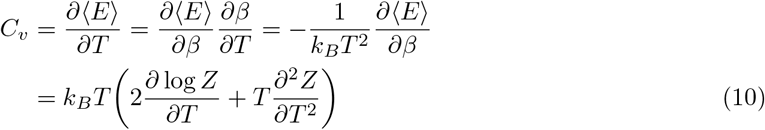

and the specific heat, which measures fluctuations in the system

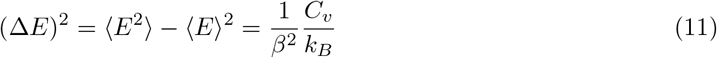

### 2.4 From Physics to Biology

Our analysis of the role of radiation and the effects of environmental factors (e.g., temperature and salinity) on the production of mutations relies on thermodynamic quantities because of the well-established associations between energy and mutations. For instance, some mutations can strengthen the binding affinity of ligands and of nucleotide base pairs, which can increase the melting temperature (i.e., resistance to perturbation) of DNA [25, 26]. Furthermore, a biophysical model based on the statistical behavior of binding sites characterizes mechanisms of mutations [27], some of which may lead to diseases [28]. Thus, we propose the three-part Hamiltonian operator described above to determine energy contributions from each part of the system and their interactions.

The Hamiltonian presented here has the feature that the interactions are clearly divided into energy contributions well separated from each other, which makes it general, in the sense that each contribution has an intuitive role in the theory. The construction of the specific terms contributing to the energy vary from author to author, for instance, in [25] energy contributions are due to van der Waals and electrostatic forces, entropy and polar and non-polar solvation, in [29] the energy is decomposed into deformation, tautomerization and stabilization, energies, taking into account the geometry of DNA and interactions inside. Our model captures the intricacies of the system in letting the parameters *h*_*i*_, *g*_*i*_ and *ϵ*_*ik*_ be free. As we will show in the next section, the determination of the free parameters characterizes the model as DNA rather than a different kind of substance. This is consistent with the difficulties of the problem of studying DNA as a quantum system, since the nature of the interactions among the system’s parts is not exactly known[30], at least to the best of our knowledge.

We point out that our model is general enough, because the thermodynamics of the system include an external environment, whose nature is determined by the frequencies *ω*_*i*_, but we also write a term of indirect effects (*g*_*i*_ determines their strength), which can be tuned to represent interactions that modify DNA without the exchange of particles. The loss of DNA-binding affinity is attributed, among other factors, to the loss of a salt bridge between the DNA chains [31], which highlights the importance of including such indirect effects in the quantum mechanical model. Therefore, our model combines relevant mutation precursors into a relatively simple description (at least intuitively).

We focus on point mutations and deletions, but we do no attempt to determine the fate of each base-pair; rather, we look for statistical effects of interactions among the DNA and radiation. We aim to characterize the set of parameters that would make mutations more likely, keeping in mind that there may be more than one set of parameters that lead to the same genomic variation. Our study is in line with the idea that genetic variation makes natural selection possible, and biased mutation rates determine variant accessibility, which may be more important to evolution than fitness [32]. Also, not knowing the fate of each pair-base is intended in our theory, taking AT as an inspiration, where selectivity is not described in terms of a specific mechanism, but it is explained if one knows the number of objects, how they can assemble, and how many of them are available [5].

## 3 Results

To have our model give relevant information about evolutionary trajectories, one should know the strength of the couplings inside the DNA, those of DNA-radiation and the strength of indirect effects beforehand, which amounts to knowing the values of the free parameters in the model. However, it is in the scope of the present manuscript to provide estimates for those values. Therefore, we take a different approach: we explore different combinations of *h*_*i*_, *g*_*i*_ and *ϵ*_*ik*_ at different temperatures to characterize their impact on the DNA-environment model.

To represent a gene in the Hamiltonian (1), we fixed *N* — the number of base-pairs — to 75; thus, our gene is composed of 25 codons. Although this synthetic gene is small in size, it is larger than the smallest known gene in nature, which consists of 21 base-pairs [33]. With this choice, there are 75 values of *h*_*i*_ and *g*_*i*_ to be determined (see eq. (5)). To represent codons, we generated a set of 20 random energy values in the interval [0 1]eV, such that the final energies are in the order of eV, as one should expect for genes. Then, in groups of three consecutive *i* values, we assigned each *h*_*i*_ the same energy from the aforementioned set, until each *h*_*i*_ has a defined value. The rationale for creating 20 energy values relies on the fact that 20 amino acids are the building blocks of proteins. In Figure 1 we illustrate the way we represented codons in the simulations.

**Figure 1.**
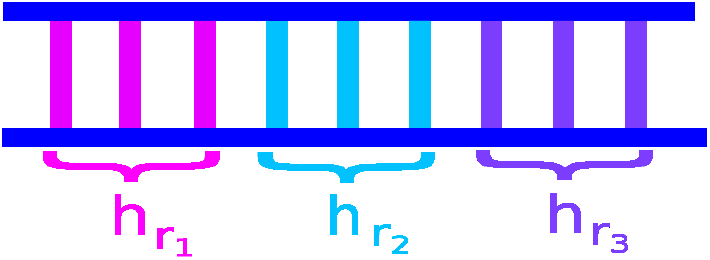
We build codons by grouping three consecutive base-pairs and assigning them the same energy. In this diagram, we show a DNA segment, containing nine base-pairs — i.e. three codons —. From a set of twenty possible energy values, we take three of them, here written as 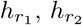 and 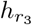, to be the energy for each codon. It is possible that two or more codons are the same, if that is the case, we give them the same energy.

To extract the thermodynamics of the interacting system,we start from the general expression of the partition function (8), whose full form is given in the appendix and then extract the energy, the heat capacity and the specific heat, as instructed in equations (9), (10) and (11), respectively. We used Mathematica 14.0 to make the calculations of the thermodynamic quantities.

We distinguish two sets of simulations, in the first, we consider the DNA interacting with monochromatic radiation, while in the second we allow more frequencies to be present in the interactions. The single-frequency model can be compared to controlled laboratory conditions, while the latter allows to study the effects under a more realistic scenario. In each case, we also divide the study into two separate cases: (1) we fix the indirect effects (*g*_*i*_ = *g* = *const*., see eq. (5)), while varying the temperature and DNA-radiation interaction strength (*ϵ*_*ik*_, see eq. (7)), and (2) we allow the indirect effects to vary as well.

In Figure 2 we show the energy and the specific heat, as the temperature and coupling strength vary. Of particular interest is the latter quantity, since it measures fluctuations in the system. As the specific heat grows, we expect the system to undergo a phase transition where its properties would change abruptly. We present findings on the energy and specific heat while holding indirect effects constant and allowing variations in temperature and DNA-radiation interaction strength, with DNA interacting with monochromatic UVC radiation. On the left panel, we illustrate how the system’s energy changes with different temperature and DNA-radiation interaction combinations. For a constant coupling, increasing temperature leads to a more negative energy, which would typically suggest stronger binding in an isolated system, but this is not the case here. On the right panel, we observe that specific heat increases with higher temperatures, also for a fixed interaction strength, indicating increased system fluctuations. These fluctuations could potentially trigger a phase transition in an infinite system, but in our case, the higher values suggest environmental influence has sufficiently perturbed the DNA to modify its behavior. Our findings underscore that mutations are influenced by both environmental factors and temperature.

**Figure 2.**
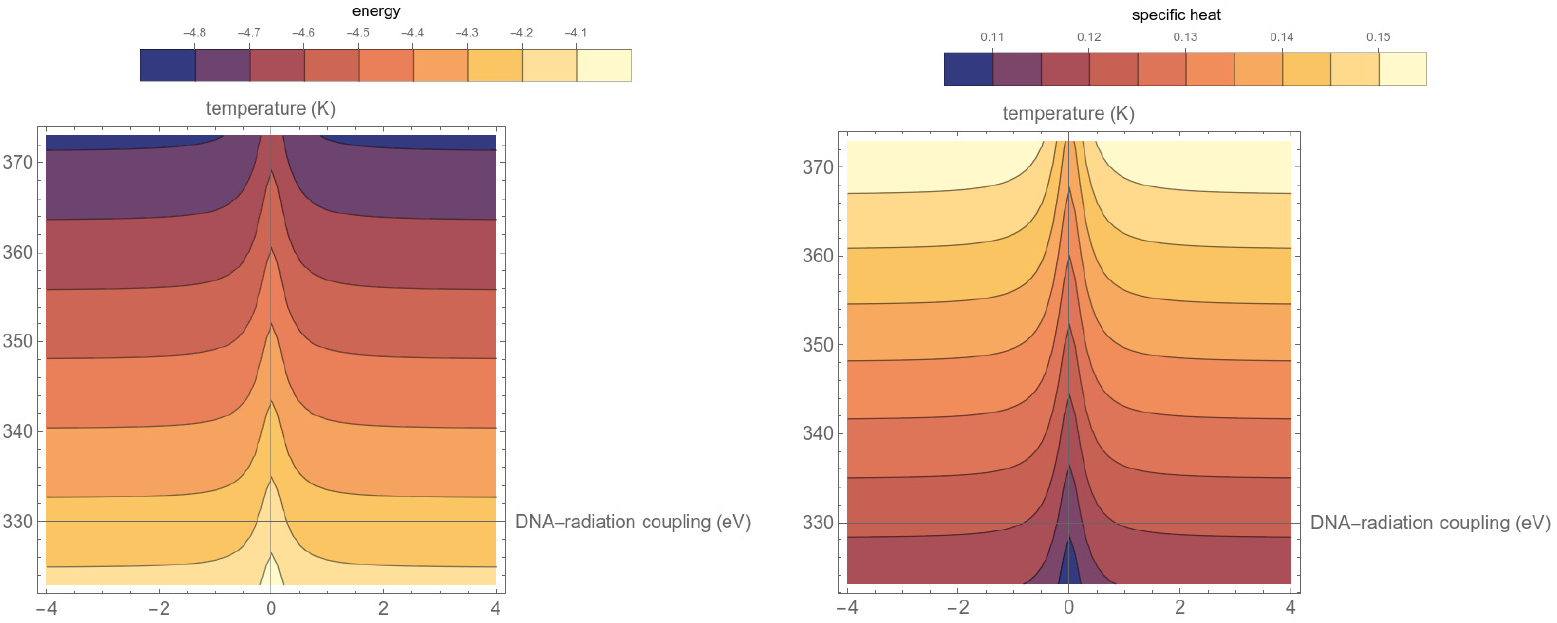
We show results for the energy and specific heat when the external effects are kept constant and let the temperature and the DNA-radiation coupling vary, letting the DNA interact with monochromatic UVC radiation. (left) We show how the energy of the system changes for different combinations of temperatures and DNA-radiation couplings. For a fixed coupling, the temperature makes the energy more negative, in an isolated system, this would mean that the whole system is more strongly bound; however, this is not the case in the present study. (right) The specific heat becomes larger at higher temperatures, for fixed coupling, thus the system tends to fluctuate more. Fluctuations in the system would eventually lead to a phase transition, in an infinite system, but here, simply by having larger values we are certain that the environment has disrupted the DNA enough, to modify its behaviour. Our results show that mutations are indeed driven both by the environment and the temperature.

We then proceeded to explore further the effects of varying the remaining free parameters of the Hamiltonian simultaneously. Now, the indirect effects and the coupling strength take on different values at different temperatures (see Figure 3). We notice that higher temperatures lead to higher specific heats, i.e. energy fluctuations are set up by increases in temperature, which characterizes critical behavior in physical systems. We allowed variations in both the indirect effects (the influence of salinity on DNA), the DNA-radiation interaction, and the temperature simultaneously, while considering a single UVC frequency as the initiator of the DNA-radiation interaction. The energy pattern exhibits similar trends to previous observations (refer to Figure 2), indicating a tighter cohesion of the system at higher temperatures. However, as depicted in the specific heat, the system’s fluctuations increase with rising temperatures (i.e., with higher radiation energy, at quantum scales the DNA base pair binding strength increases amid stronger fluctuations in the system due to higher specific heat). Both energy and specific heat exhibit symmetric patterns, attributed to our second-order approximation of the partition function, implying that the interaction strength and indirect effects have comparable impacts on the system’s overall behavior. However, this scenario changes when multiple UVC radiation frequencies are introduced (see below).

**Figure 3.**
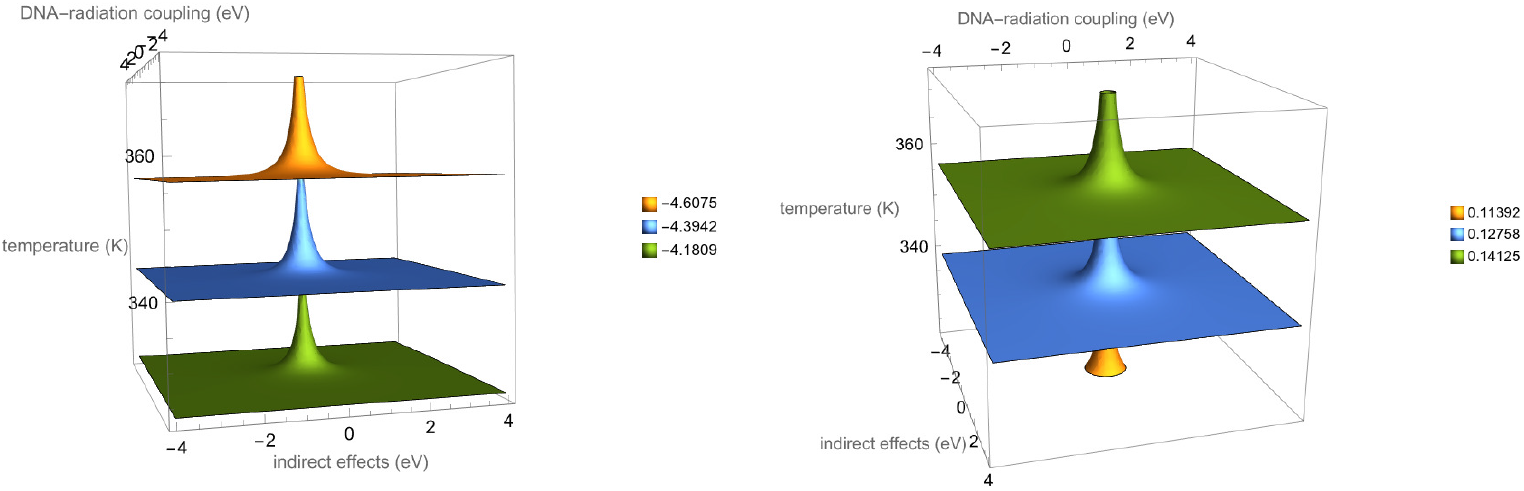
We then let both the indirect effects (the role of salinity on DNA), the DNA-radiation coupling and the temperature vary simultaneously, considering a single UVC frequency as the source of the DNA-radiation interaction. (left) The energy follows the same qualitative behaviour as before (see Figure 2), making the system appear more tightly bound for larger temperatures, but as (right) the specific heat shows, the fluctuations in the system are larger for larger temperatures. Notice that both the energy and the specific heat have symmetric shapes, this is due to the fact that we have made a second-order approximation to the partition function, thus the coupling strength and the indirect effects have similar effects on the overall behavior of the system. This changes when more than one frequency is present.

### 3.1 Three bosonic modes

To explore a more natural scenario, our model allows us to input more than one UVC frequency. Due to computing power restrictions, we made *M* = 3, which is representative of natural light composed of three frequencies, each with the capacity to alter the DNA. Here, we did the same as in the previous section, we chose three random frequencies, such that *ω*_*i*_ ∈ [4, 12] eV and kept *g*_*j*_ = *g* for *j* ∈ {1, *N* }, and measured the thermodynamic quantities; then, we also let each *g*_*j*_ vary and also measured the quantities of interest.

As one can see in Figures 4 and 5, the qualitative behaviour follows the same pattern as in the single bosonic mode case; however, now the specific heat requires a weaker coupling of the systems to grow, and the temperature effects are seen earlier. In the case where all free parameters are allowed to vary, we notice that the geometric symmetry of the heat capacity and the specific heat is broken, as expected, since now the radiation effects dominate the indirect effects.

**Figure 4.**
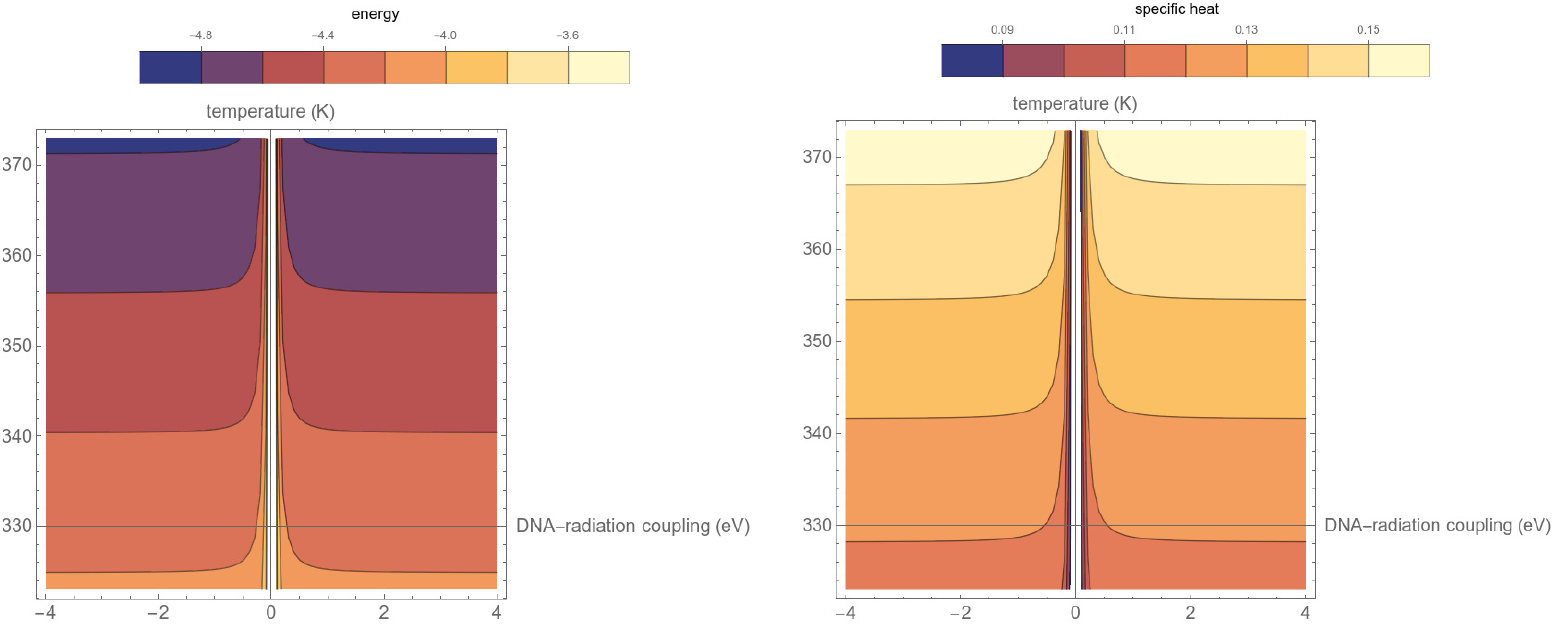
We let three UVC radiation frequencies interact with our gene. First, we fix the indirect effects, letting the DNA-radiation coupling and temperature vary. The qualitative behaviour of both the energy (left) and the (right) specific heat follow the same pattern as those shown in Figure 2, but as we would expect, the energies are larger than in the previous case due to the presence of high-energy photons and, more importantly, the fluctuations are also larger than in the previous case. This shows that the system is more susceptible to changes as more frequencies are added to the system, which is what one would see in a realistic biological scenario.

**Figure 5.**
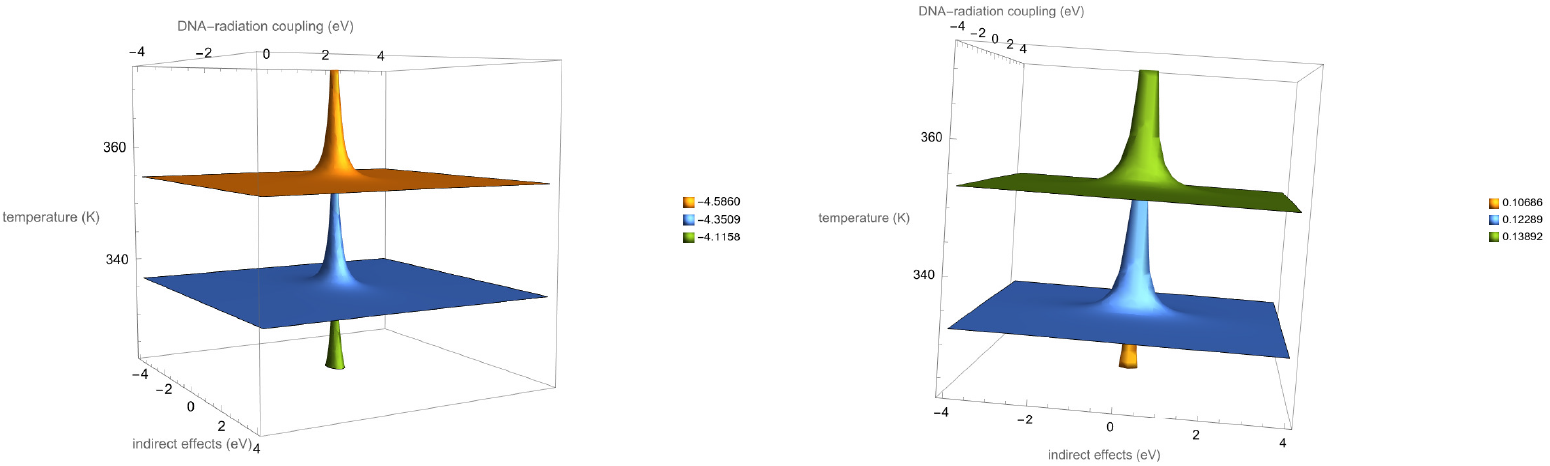
The (left) energy and (right) specific heat when the indirect effects, DNA-radiation coupling and temperature vary simultaneously. Qualitatively, these results resemble those shown in Figure 3, but we notice that the symmetry induced in the results by the second-order approximation to the partition function is now gone. Here, we see that the DNA-radiation coupling is more important than indirect effects and temperature alone, because of the highly-energetic nature of the radiation under study. The more radiation present, the faster the mutations take place.

In Figure 4, we present the results of exposing our gene to three different UVC radiation frequencies. Initially, we kept the indirect effects constant while allowing variations in DNA-radiation coupling and temperature. The qualitative trends observed in both energy (left) and specific heat (right) mirror those depicted in Figure 2. However, the energies are higher compared to the previous scenario due to the presence of more high-energy photons, and notably, the fluctuations (measured by the specific heat, on the right panel) are also amplified. This indicates a heightened susceptibility to mutations as additional radiation frequencies are introduced, reflecting a more realistic biological scenario.

In Figure 5, the energy (left) and specific heat (right) are observed as the indirect effects, whereas DNA-radiation coupling and temperature are modified simultaneously. These plots bear qualitative similarity to those illustrated in Figure 3; however, we observe that the symmetry previously introduced by the second-order approximation to the partition function is absent here. This suggests that DNA-radiation coupling holds greater significance compared to indirect effects and temperature alone, owing to the highly energetic nature of the radiation under consideration. As the radiation intensity increases, the pace of mutations accelerates.

The choices of values for the coupling constants lead to consistent results, in the sense that the energy of the system is of the order of electron-volts, as one should expect in the case of molecular systems. The energy becomes more negative at higher temperatures, which one could mistake as indicating a more strongly-bounded molecule; however, when seen in conjunction with the specific heat, we notice that the system is less stable.

From our results, we see that the specific heat shows more fluctuations as the temperature increases and the temperature is the only relevant variable when all couplings go to zero. In the interesting case, where the couplings are non-zero, their effect is to reduce the temperature at which the composite system suffers more fluctuations. This is in agreement with the behaviour of the heat capacity, whose geometric symmetry is also broken when letting all free parameters vary and more frequencies are present to disturb the DNA.

## 4 Biological interpretation

The Hamiltonian in this work is general, because it considers the interaction between any two-level system (i.e. a gene being expressed or not) and a bosonic bath (UVC radiation). We turn it into a model of DNA-radiation interaction by creating artificial codons, determined by the base-pairs binding energies, and using the Pauli operators to represent DNA.

There are two interaction terms, one quantifies the direct effects of radiation on the stability of the DNA molecule, *ϵ*_*ij*_, while another accounts for indirect effects, *g*_*i*_, such as salinity (see equations (7) and (5)). The interpretation of *ϵ*_*ij*_ is the following: a stronger interaction implies that radiation is more likely to modify the DNA chain. Moreover, using the parameters *g*_*i*_ allows us to account for indirect effects on the DNA without having to define the mechanistic details (i.e. all-on-all interactions) of the system.

We make educated guesses of the values of interaction strength to determine their effects on a realistic DNA system. Our results offer consistency to biological data and the possibility to further explore our quantum statistical method. Firstly, the digital simulations of our model produce system energies within the order of electron-volts - hence, in agreement with experimental results - thus providing evidence that the choice of parameter combinations is biologically correct. Secondly, higher temperatures lead to more fluctuations of the system, regardless of the interaction strengths. Finally, when the interactions are present, our results show that the system would undergo an unstable behavior at lower temperatures. This critical behaviour might ultimately be connected to DNA denaturation, but even in the absence of denaturation, the fact that the specific heat (a measure of energy fluctuations) is larger at lower temperatures means that DNA energy fluctuations are driven by the environment, via direct and indirect effects.

Providing experimental data to the model will help us establish values for the interaction strengths and, more importantly, we will be able to determine a normalized specific heat value that characterizes a gene’s mutations across the different experimental adaptation stages. To test our model, we have simulated it in digital computers. Next steps will include running our model in analog quantum computers capable of performing quantum-quantum simulation or, alternatively, transforming our quantum walks-based model into quantum circuits.

## 5 Conclusions and outlook

In this manuscript we have presented a mathematical model relying on quantum statistical mechanics to study point mutations. We provided evidence that determining the thermodynamic properties of the system results in energy scales comparable to those of realistic DNA models. Furthermore, the measurement of fluctuations in the system also characterizes how the temperature, radiation, and indirect effects (in this case, salinity) combine to drive the dynamics of the open quantum system.

Our model is novel, since it incorporates both direct (radiation) and indirect (salinity) effects into terms that capture the energetic changes inside the DNA molecule. Also, this makes our model general, since we do not need to know specific mechanisms in the system, we just need to know their effect on the molecule. Due to its statistical nature, our method allows convergence to the same outcome (i.e., a mutation occurring or not) regardless of the base-pair composition given that the dynamics are determined by the system’s total energy.

For the time being, the free parameters in the model are set by requiring that the energetic scale is of the order of electron-volts. However, we aim to use experimental data (1) to refine the values of those parameters and (2) to compare the specific heat values at different salinities and radiation conditions, in order to establish a one-to-one relation between real DNA evolution and numbers resulting from our model. We are currently analyzing genomic information derived from adaptive laboratory evolution experiments with algae.

## A Model Dynamics: Mathematical Methods

To compute the trace in eq. (8) we need to sum over all possible configurations of the interacting system, that is, we have to sum over the DNA and the bosonic states. Since we do not know the eigenvalues of the Hamiltonian, this is a challenging task. A possible strategy to approximate the exponential in the partition function uses the Zassenhaus formula [34],

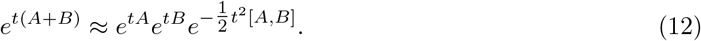

where *A* and *B* are non-commuting operators and *t* a parameter that corresponds to *β*, in our case. However, the approximation may be non-Hermitian [35].

Instead of using the previous approach, we considered the Hermitian approximation presented in ref. [23]

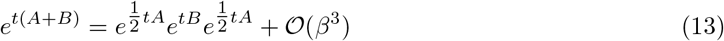

where, due to the cyclic property of the trace,

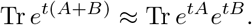

We have chosen *A* = *H*_*bath*_ and *B* = *H*_*DNA*_ + *H*_*interaction*_ to carry our calculation of the partition function as

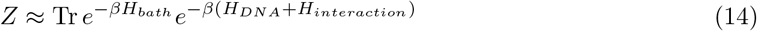

we also notice that

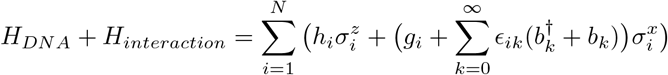

In the following section, we present our calculations leading to an expression of the partition function that we can work with.

### A.1 Calculation of the Partition Function

We start by taking the trace over the DNA variables, (in the following, Tr _*b*_ represents the trace operator over the bosonic bath), keeping in mind that we trace ove the Pauli operators, such that the eigenvalues of *σ*^*z*^ are ±1,

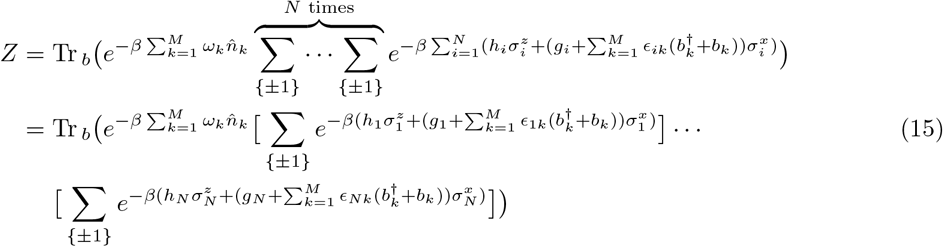

We can expand each exponential appearing the second equality as

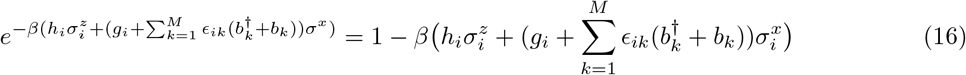

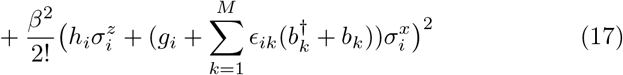

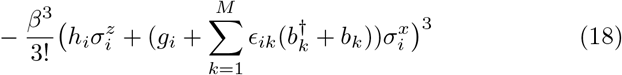

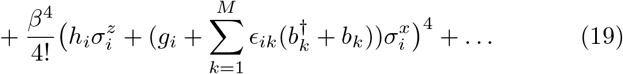

It is convenient to write

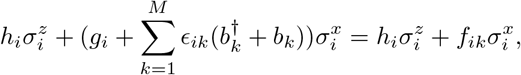

with

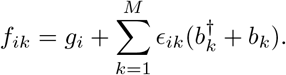

If we work explicitly the quadratic term, 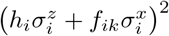, we see that

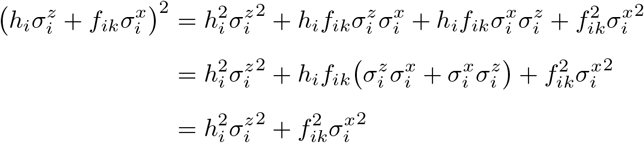

where we have used that 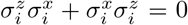. It is then striaghtforward to generalize, and notice that for *k* ≥ 2, each term in the series can be written as

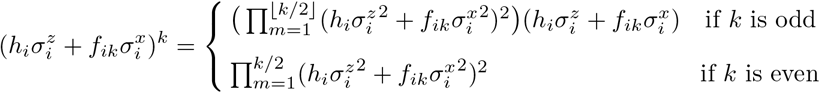

Then, we can write the exponential (16) as

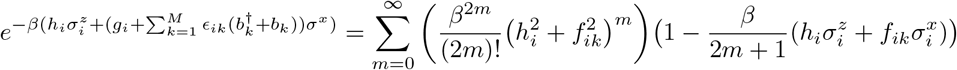

Since the Pauli matrices are traceless, after carrying out the sums over spin states, the partition function (15) becomes

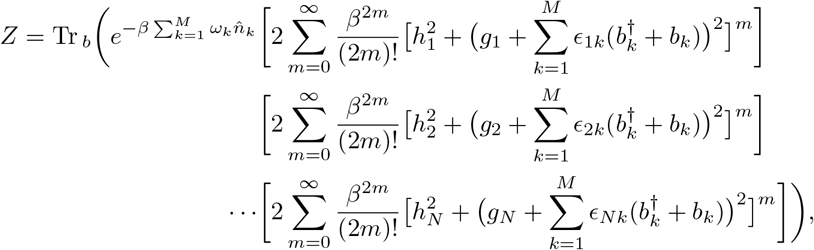

that is

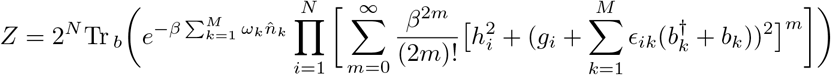

This completes the tracing over the DNA variables.

We now have to take the trace over the bosonic fields. To do so, we employ the Fock states for each harmonic oscillator. To carry out that calculation, it is useful to notice that equations (4) imply that

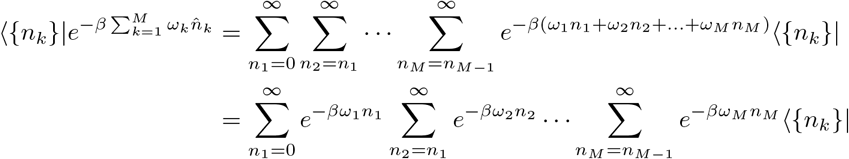

from which we get

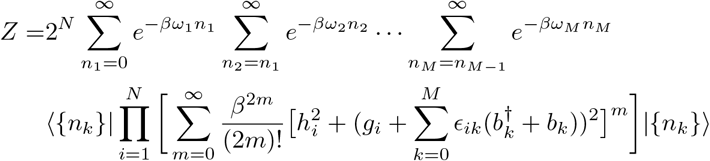

In addition, we see that

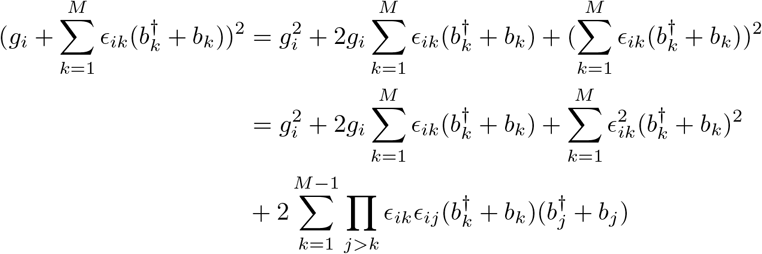

Since we are interested in the cases where *β* is small (high temperatures), we keep terms of order *β*^2^ our calculation. By combining our two previous results, we get

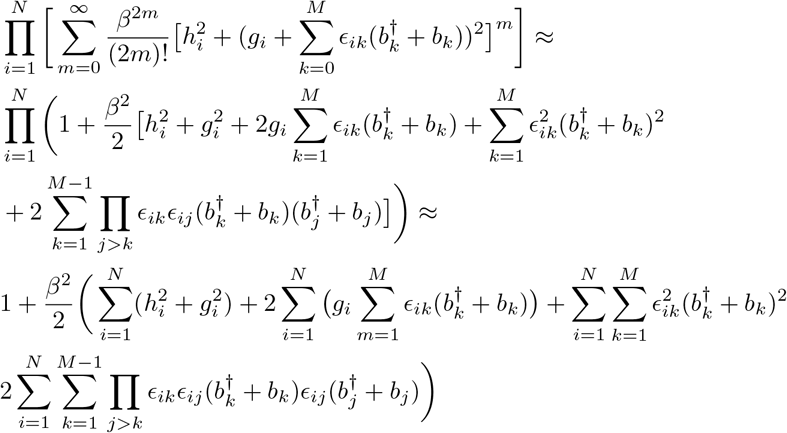

Because of the action of the creation and annihilation operators on the number states (see eq. (4)), the linear terms, 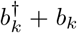, in the previous equation will not contribute to the trace; the same holds true for powers of 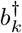 and *b*_*d*_. From the commutation relations (2), we have that

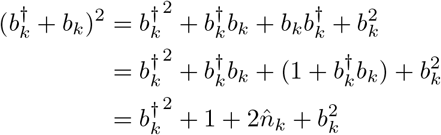

The considerations above allow us to approximate the partition function as

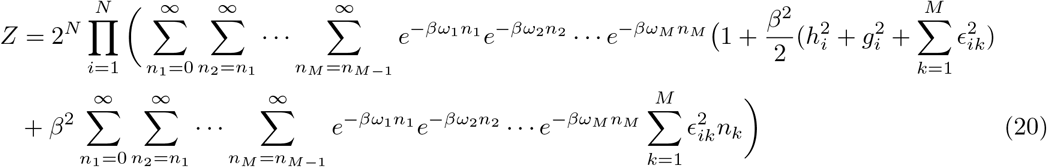

and the next task is to evaluate the summations over *n*_*k*_. We start with the first term on the r.h.s. which corresponds to *M* uncoupled oscillators. Since

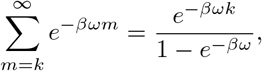

Then

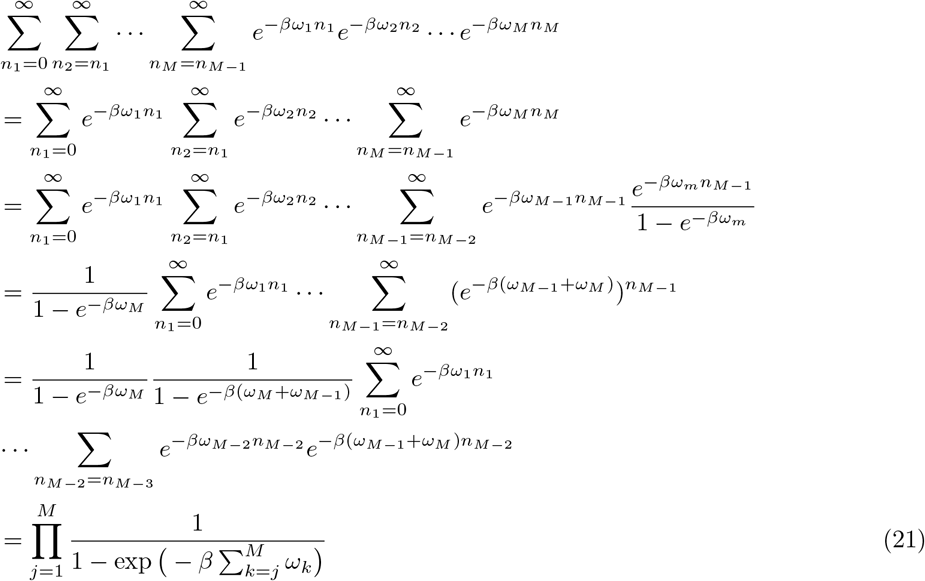

We go back to equation (20), noticing that

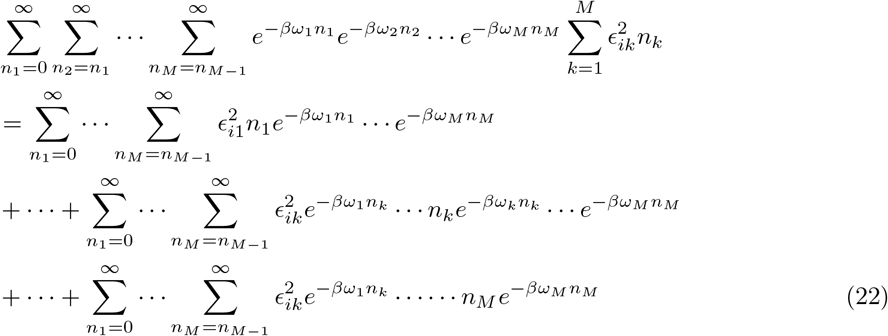

Let us work out a general term of the equation above, using the same procedure that lead us to equation (21), for *k < M*,

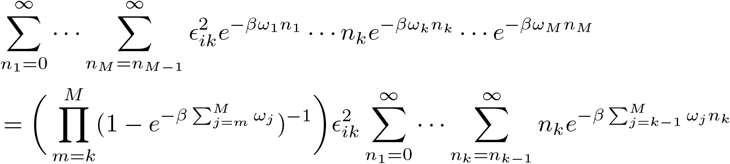

Let us evaluate the final summation in the previous equation,

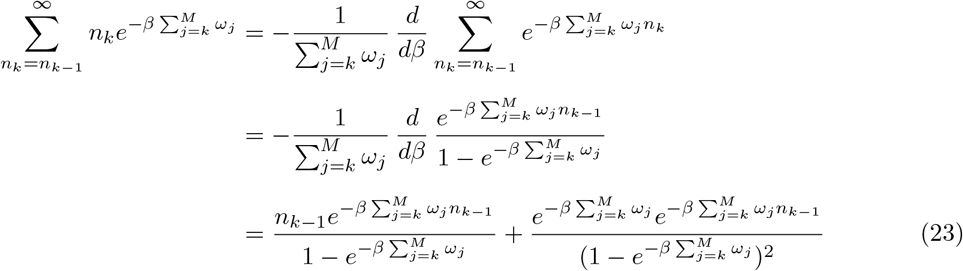

Upon substitution of (23) in (22)

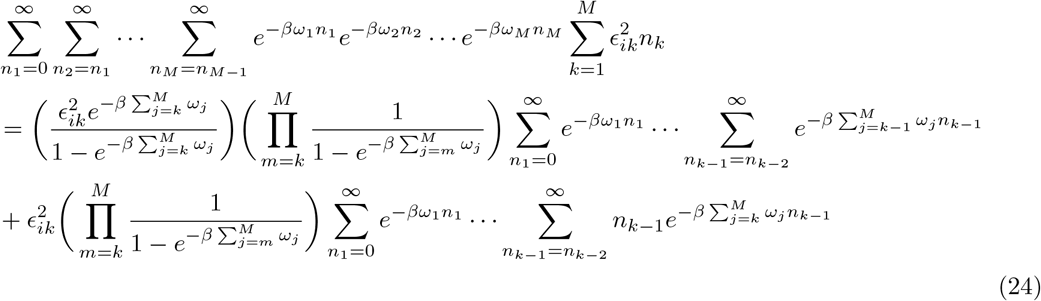

The first term on the right-hand-side of the equation above yields the result

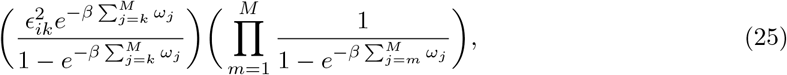

which is similar to that of the uncoupled oscillators term, multiplied by a constant term. Let us notice that when *k* = *M*, the total result will resemble what we get from the second term.

The second term in eq. (24) induces a recursivity relation, where summations of the form 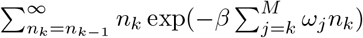 keep on appearing; we proceed as we have before to calculate all the summations.

After calculating all of the summations, we end up — for *k < M* — with

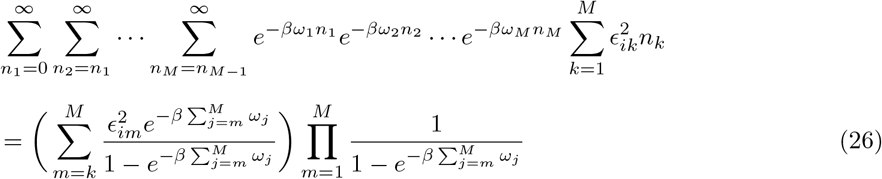

We then have to repeat this process for each possible value of *k*, including *k* = *M*, which we have omitted so far. However, all terms will be of the same form; we will have

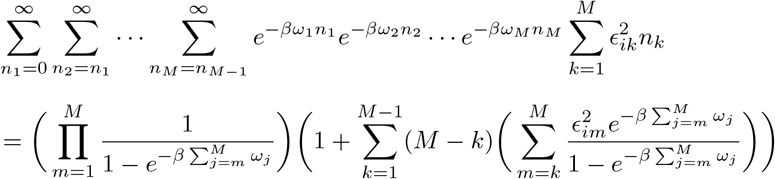

Finally, we plug this result in the partition function, eq. (20) to get

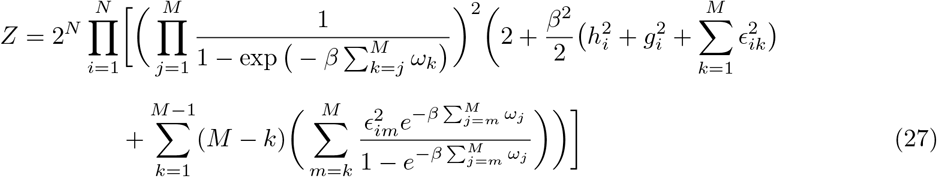

## Acknowledgements

HM-D acknowledges the financial support by CONAHCyT [CVU 560425]. DS-A acknowledges the financial support of USF via startup funds. SEVA acknowledges the financial support provided by Tecnologico de Monterrey, Escuela de Ingenieria y Ciencias and CONAHCyT-SNI [SNI number 41594]. SEVA acknowledges the unconditional support of his family.

## Footnotes

1 Under most circumstances, the Hamiltonian operator is the operator from which we can extract the possible energy values of a given system.

2 Since *σ*^*y*^ is not involved in our model, we omit it from here onward. For a detailed discussion of the Pauli operators see, for example [19].

3 The only operator that we represent with a *hat* is the number operator, 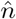, where confusion with its eigenvalues, *n*, may occur.

4 We have set *ω* = *ħω*_*freq*_, where *ω*_*freq*_ is the frequency of the radiation and *ħ* is the reduced Planck’s constant.

